# SLAYER: A Computational Framework for Identifying Synthetic Lethal Interactions through Integrated Analysis of Cancer Dependencies

**DOI:** 10.1101/2024.05.01.592073

**Authors:** Ziv Cohen, Ekaterina Petrenko, Alma Sophia Barisaac, Enas R. Abu-Zhayia, Chen Yanovich-Ben-Uriel, Nabieh Ayoub, Dvir Aran

## Abstract

Synthetic lethality represents a promising therapeutic approach in precision oncology, yet systematic identification of clinically relevant synthetic lethal interactions remains challenging. Here we present SLAYER (Synthetic Lethality AnalYsis for Enhanced taRgeted therapy), a computational framework that integrates cancer genomic data and genome-wide CRISPR knockout screens to identify potential synthetic lethal interactions. SLAYER employs parallel analytical approaches examining both direct mutation-dependency associations and pathway-mediated relationships across 808 cancer cell lines. Our integrative method identified 4,332 statistically significant interactions, which were refined to 142 high-confidence candidates through stringent filtering for effect size, druggability, and clinical prevalence. Systematic validation against protein interaction databases revealed a 15-fold enrichment of known associations among SLAYER predictions compared to random gene pairs. Through pathway-level analysis, we identified inhibition of the aryl hydrocarbon receptor (AhR) as potentially synthetically lethal with RB1 mutations in bladder cancer. Experimental studies demonstrated selective sensitivity to AhR inhibition in RB1-mutant versus wild-type bladder cancer cells, which probably operates through indirect pathway-mediated mechanisms rather than direct genetic interaction. In summary, by integrating mutation profiles, gene dependencies, and pathway relationships, our approach provides a resource for investigating genetic vulnerabilities across cancer types.

## Introduction

Synthetic lethality represents a promising therapeutic strategy in precision oncology by exploiting genetic vulnerabilities exclusive to tumor cells [1]. This approach targets non-oncogenic partners of mutated genes that cancers become dependent on, providing a means to selectively kill cancer cells while sparing normal tissues [2]. Notable examples of clinically relevant synthetic lethal interactions include PARP inhibitors in BRCA-mutated cancers [3] and WEE1 inhibitors in TP53-mutated tumors [4].

The systematic identification of synthetic lethal interactions (SLIs) has been pursued through various computational approaches. Early methods focused on evolutionary conservation of genetic interactions [5], while more recent approaches have leveraged cancer genomic data. For instance, computational frameworks like DAISY integrated somatic copy number alterations, gene expression, and shRNA data to predict SLIs [6]. Another approach, MiSL, analyzed mutation and copy number profiles to identify candidate synthetic lethal partners [1]. The advent of genome-wide CRISPR screens has enabled more direct approaches, as demonstrated by Chan et al. in identifying WRN as a synthetic lethal target in microsatellite unstable cancers [7]. Recent studies have highlighted the importance of pathway-level analysis in SLI prediction. For example, the ISLE framework incorporated pathway information to prioritize SLI candidates [1], while others have focused on specific pathway contexts, like DNA damage response genes [8]. However, most existing approaches either rely on direct gene-gene associations or treat pathways as discrete sets, potentially missing complex regulatory relationships that could influence synthetic lethality.

The Cancer Dependency Map (DepMap, https://depmap.org/portal/) provides comprehensive multiomics data spanning over 1,000 cancer cell lines, including genome-wide CRISPR-Cas9 knockout screens that measure cellular viability upon individual gene disruptions. While this resource has been leveraged for various analyses, systematic integration of mutation profiles, gene dependencies, and pathway-level information remains challenging. Previous studies using DepMap data have primarily focused on direct mutation-dependency associations through large-scale CRISPR screens [9] of pathway-level information.

Here, we present SLAYER (Synthetic Lethality AnalYsis for Enhanced taRgeted therapy), a computational framework that integrates multi-dimensional DepMap data to systematically identify associations between common cancer mutations and pathway dependencies. SLAYER builds upon previous approaches in three key ways: (1) it combines direct mutation-dependency analysis with pathway-level investigation, (2) it incorporates druggability and clinical relevance filters to prioritize actionable targets, and (3) it enables exploration of indirect synthetic lethal relationships mediated through pathway rewiring. By intersecting mutation profiles with CRISPR gene essentiality patterns and pathway enrichment scores, our approach provides a systematic framework for nominating potential SLIs for experimental validation and therapeutic development.

## Results

The SLAYER computational framework integrates multi-dimensional data from the DepMap resource to systematically identify SLIs across cancer types. Our analysis encompassed 808 cancer cell lines with complete genomic and transcriptomic profiles alongside CRISPR-Cas9 gene essentiality screens. Initial mutation profiling identified 120 commonly mutated genes based on a 5% recurrence threshold across the cell line panel spanning 31 cancer types.

Using the DepMap CRISPR-Cas9 knockout viability data, we calculated gene-specific dependency scores and their associations with driver gene mutations to identify potential SLIs through complementary analytical approaches (Figure 1). The primary analysis examined direct associations between mutations and gene dependencies, while a parallel pathway-oriented analysis investigated potential indirect relationships by leveraging experimentally-derived oncogenic signatures from MSigDB to capture broader cellular context (Methods). This integrated analysis yielded 4,332 statistically significant potential SLIs (adjusted p-value ¡ 0.05) with delta dependency scores exceeding 0.2 across both analytical approaches.

**Figure 1.**
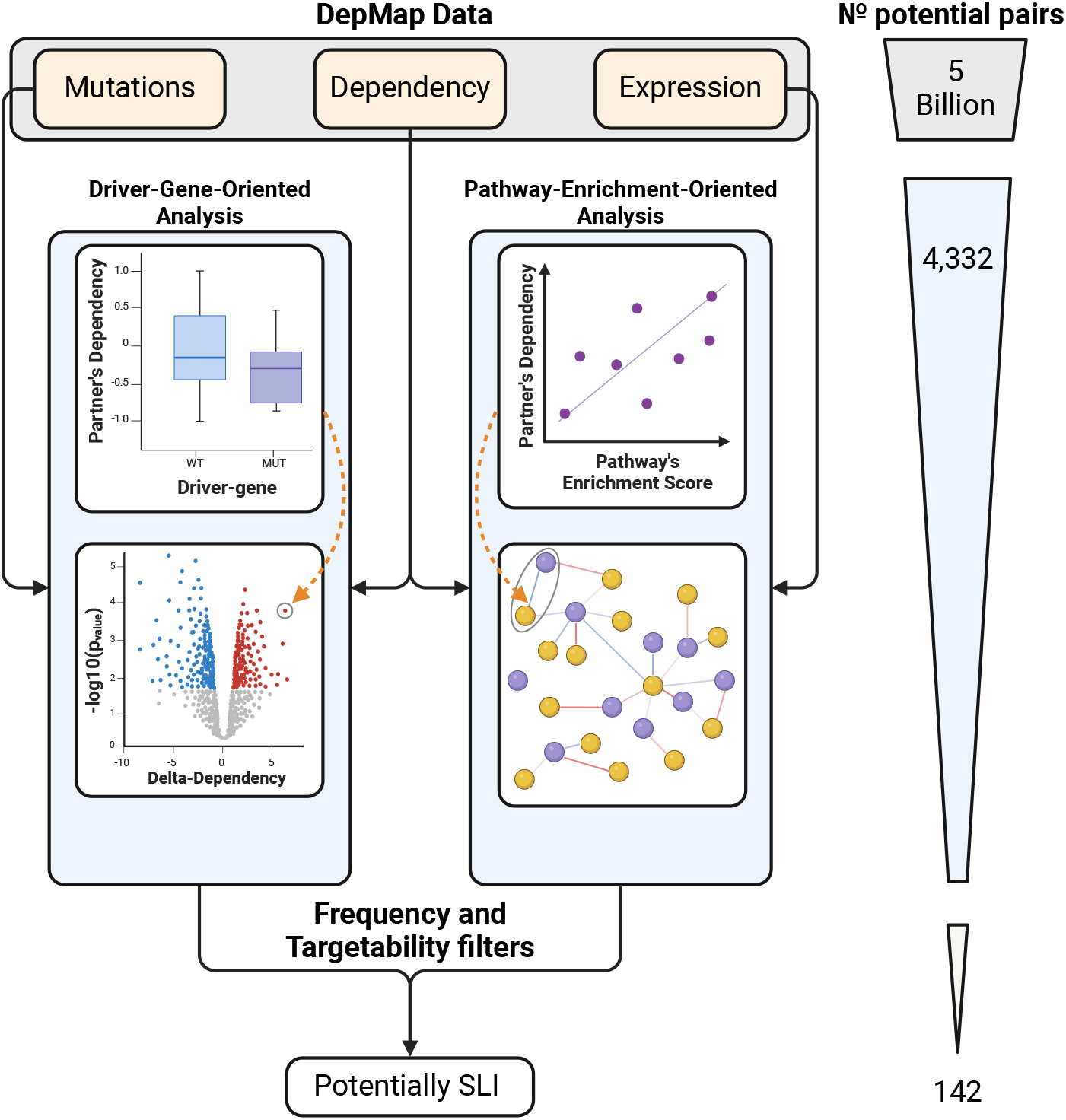
Schematic overview of the SLAYER methodology. Data from the DepMap portal, including mutations, gene dependencies, and expression profiles across 808 cancer cell lines, were integrated using two complementary approaches: (i) a driver gene-oriented analysis associating gene dependencies with driver mutations, and (ii) a pathway enrichment-oriented analysis connecting gene dependencies with mutation-associated pathway activities. Putative SLIs were identified from each approach and further filtered based on the driver gene mutation frequency in the patient population and druggability of the partner gene. From an initial 5 billion possible gene pairs, 4,332 statistically significant SLI candidates were identified, which were then refined to a set of 142 high-confidence, actionable SLIs based on effect size, targetability, and clinical relevance filters.

We then applied stringent filtering criteria to prioritize clinically actionable candidates. First, we filtered for partner genes that are possible therapeutic targets by cross-referencing against the DepMap PRISM repurposing data, retaining only SLIs where the partner gene is inhibitable by an existing drug or compound. Next, we filtered for SLIs involving driver gene mutations with a population frequency greater than 15% based on TCGA data in cBioPortal. This ensures the SLIs are applicable to a sufficiently broad patient population. After this multi-step filtering process focused on statistical significance, effect size, druggability, and clinical relevance, we identified a refined set of 142 high-confidence, actionable potential SLIs for further prioritization and experimental validation (Figure 2 and Supplementary Table 1).

**Figure 2.**
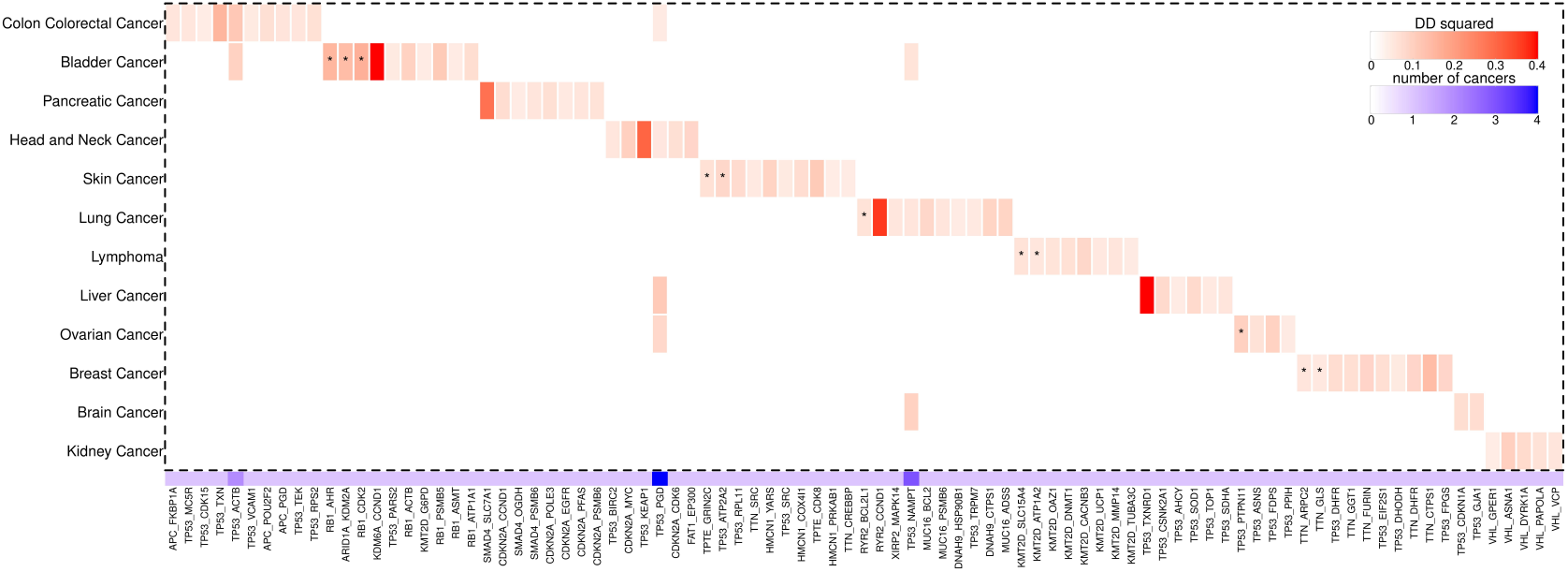
Potential synthetic lethal interactions across cancer types. The heatmap displays the top 10 most significant SLIs identified for each cancer type in the analysis. The red color scale corresponds to the delta dependency (DD) score, representing the difference in gene essentiality between the mutant and wildtype groups. The blue bar at the bottom indicates the number of cancer types where the SLI was determined as statistically significant. SLIs with a *p* − *value* < 0.001, are marked with an asterisk.

Analysis of our predictions against STRING-db, a comprehensive database integrating known and predicted protein-protein interactions, revealed remarkable concordance with known biology (Table 1). Ten of the top fourteen most significant synthetic lethal interactions exhibited evidence of gene pair co-occurrence or interaction, including the STK11-ATP1A1 interaction in lung cancer previously validated as synthetically lethal in vivo [10], and the RYR2-BCL2L1 relationship implicated through calcium signaling studies [11]. In addition, noteworthy among the top pairs was the SMARCA4-EP300 interaction in lung cancer, which has been previously suggested in the context of synthetic lethality through independent experimental studies [12, 13]. This validation rate of 71.4% among top predictions significantly exceeded background expectations. When examining the complete set of 142 high-confidence predictions, 26.1% showed STRING-db support compared to only 1.0-5.8% for random pairs of driver genes and partner genes (Figure 3). This striking 15-fold enrichment highlights the ability of our integrated analytical approach to capture biologically meaningful relationships.

**Table 1:**
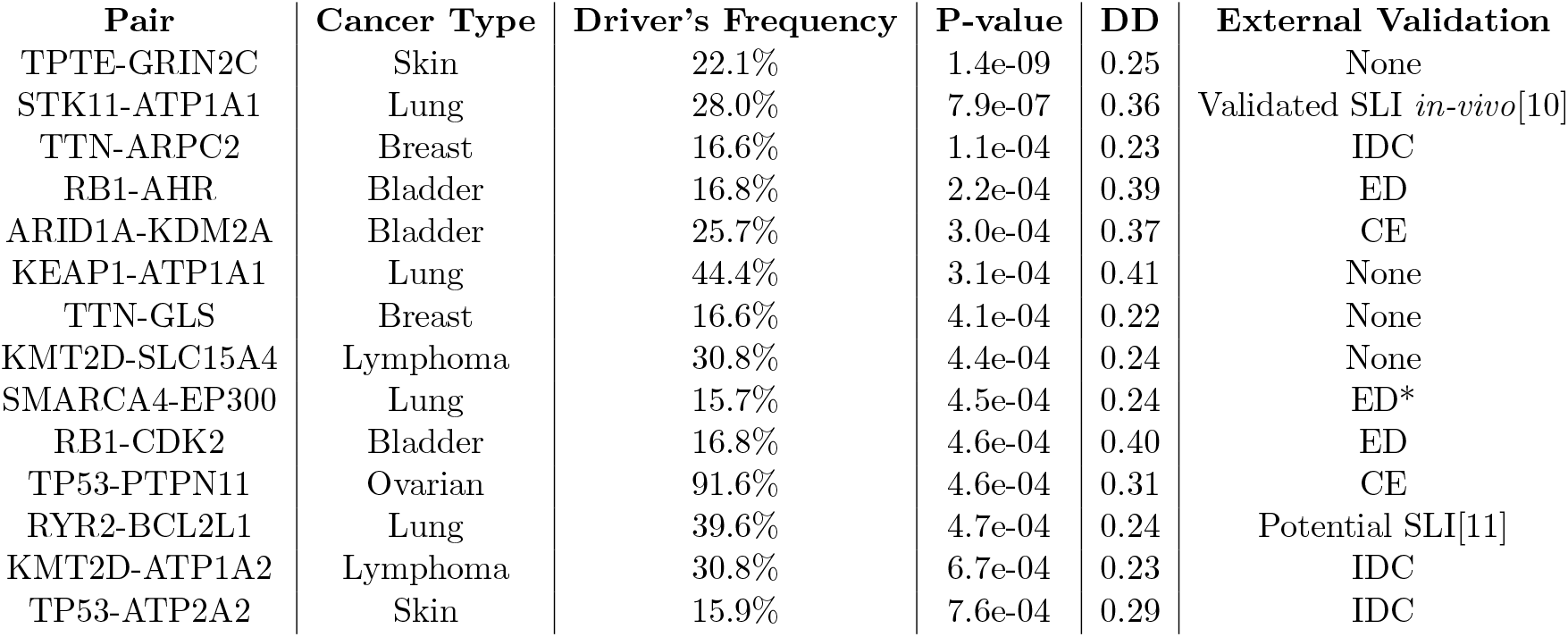
Top potential synthetic lethal interactions. ED - Experimentally Determined, CE - co-expression, IDC - Indirect connection. * Co-mentioned in the context of SLIs [12, 13].

**Figure 3.**
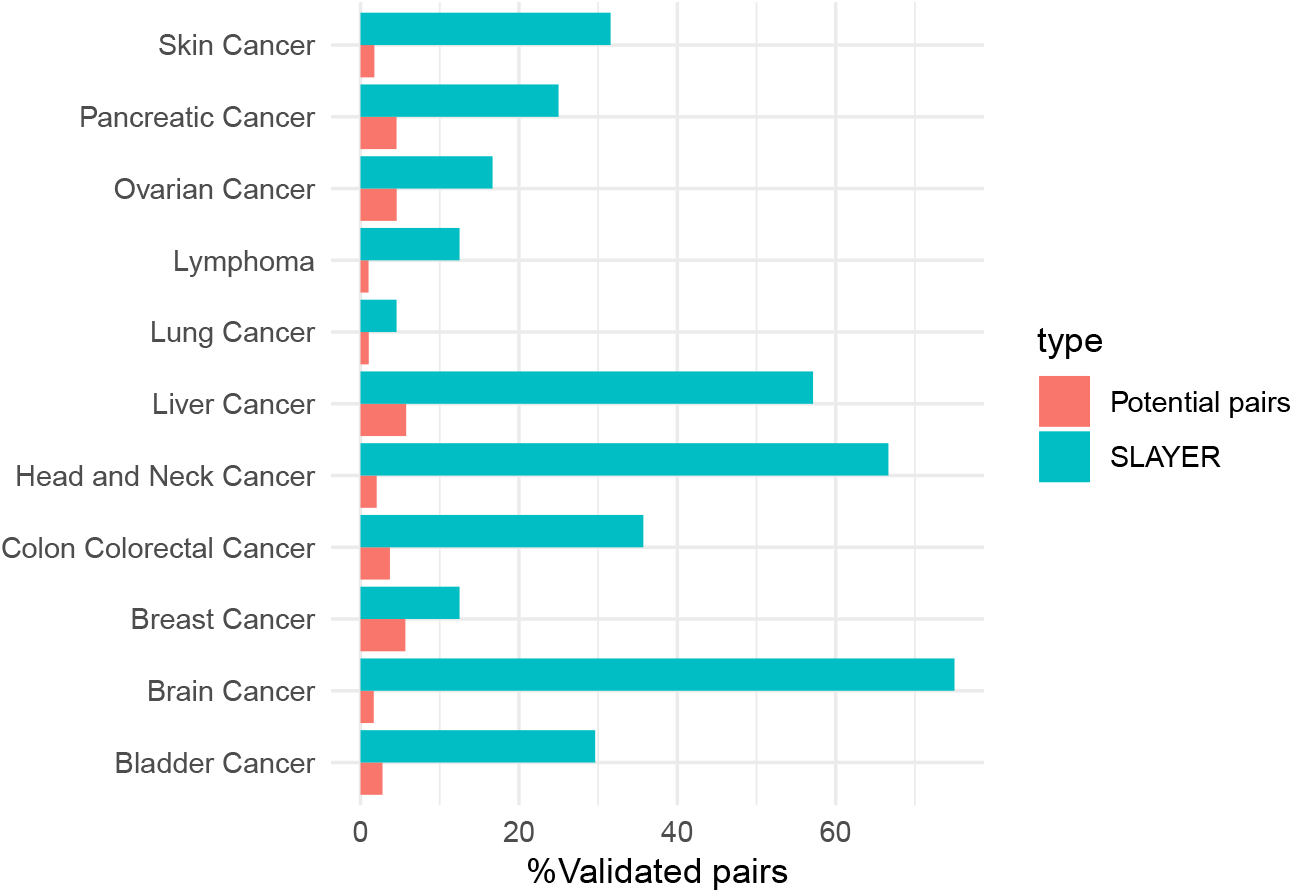
Validation of predicted synthetic lethal interactions against STRING-db database. Bar plots show the percentage of gene pairs from the SLAYER analysis that exhibited evidence of interaction or co-occurrence in the STRING-db database, which integrates known and predicted protein associations from multiple sources. The blue bars represent the percentages for the 142 high-confidence SLIs identified by SLAYER, while the red bars indicate the background percentages when analyzing all possible pairwise combinations of driver genes and genes across the different cancer types.

In bladder cancer, our analysis focused on 46 cell lines harboring 7 recurrent driver mutations (Figure 4A; Similar analyses for other cancer types are available in Supplementary Figures 1-2). The top SLIs implicated ARID1A mutant sensitivity to KDM2A knockout and two potential vulnerabilities in RB1 mutant tumors through synthetic lethality with AHR and CDK2 inhibition (Figure 4B-C). RB1 mutations were observed in 18.9% of the bladder cancer cohort, consistent with 16.8% prevalence in TCGA patient samples. Pathway enrichment analysis confirmed associations between AHR knockout dependency and RB1-related signatures, prompting experimental validation of this SLI (Figure 4D-E).

**Figure 4.**
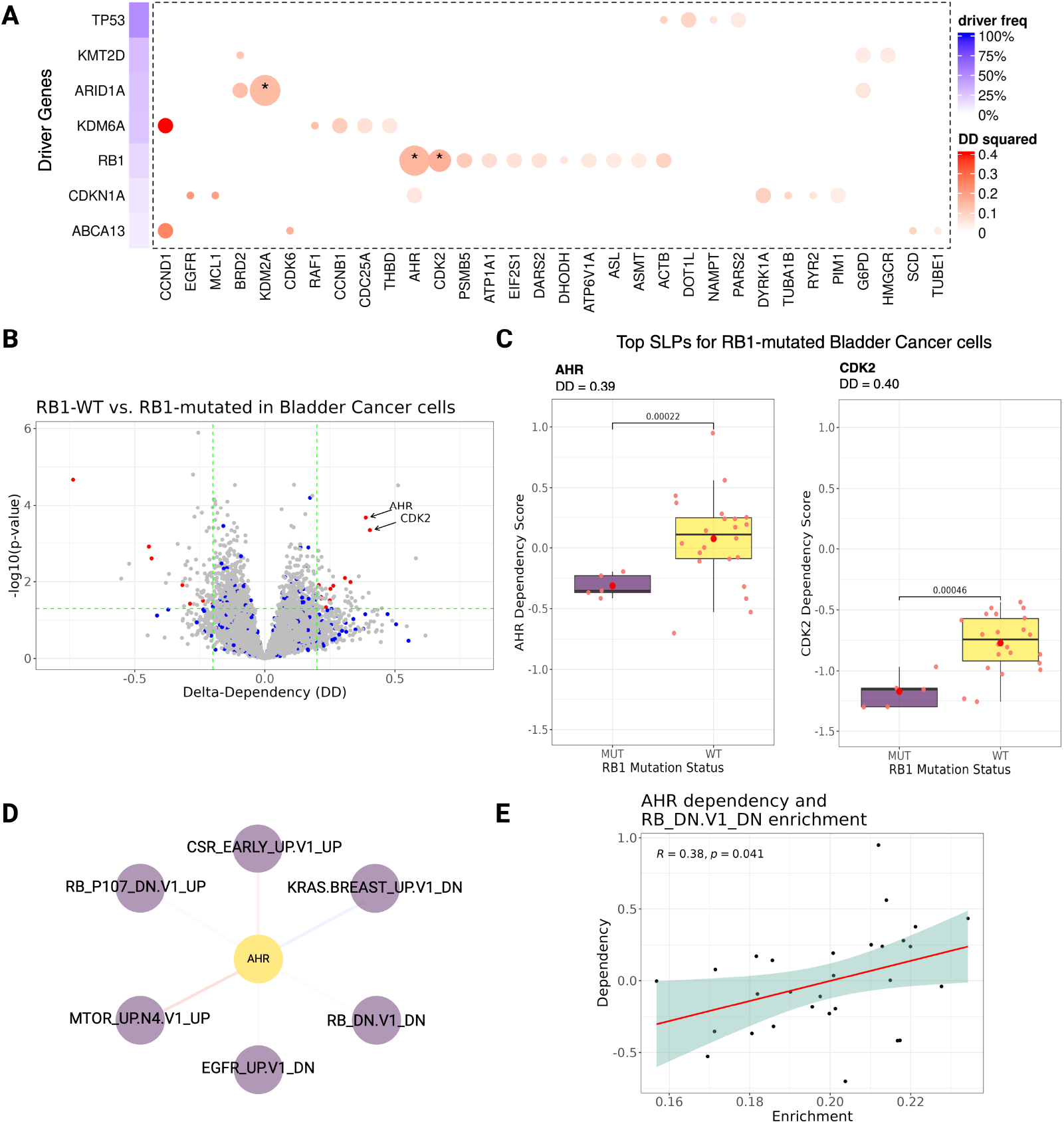
Potential synthetic lethal interactions in bladder cancer. **A**. Heatmap depicting the top synthetic lethal interaction (SLI) pairs in bladder cancer, with redder colors indicating higher delta dependency scores. Similar heatmaps for each of the cancer types are available in Supplementary Figure 1. **B**. Volcano plot showing all potential synthetic lethal interactions for the RB1 mutation, with AHR and CDK2 identified as the top-ranking candidates based on delta dependency (x-axis) and statistical significance (-log10 p-value on y-axis). More examples in Supplementary Figure 2. **C**. Boxplots illustrating the separation in gene dependency scores between RB1 wild-type and RB1 mutant bladder cancer cell lines for the AHR and CDK2. **D**. Dependency network for AHR in bladder cancer, highlighting edges that connect these genes to RB1-associated pathways. **E**. Correlation plot between the dependency scores of AHR and the enrichment of RB1-related pathway across the cell lines.

To provide preliminary experimental support for this computational prediction, we examined cellular responses to the AHR inhibitor BAY-2416964 in bladder cancer cell lines. The RB1-mutant cell line 5637 exhibited markedly increased sensitivity to AHR inhibition compared to the RB1-wildtype T24 line (Figure 5A). Importantly, BAY-2416964 treatment showed no significant impact on non-cancerous RPE cells, supporting the context-specific nature of this synthetic lethal interaction (Figure 5B). We then tested whether acute RB1 loss would sensitize cells to AHR inhibition by depleting RB1 in T24 cells using shRNA. Despite achieving significant RB1 knockdown, RB1-depleted T24 cells showed no enhanced sensitivity to BAY-2416964 (Figure 5C-D). This suggests the synthetic lethality may arise through indirect mechanisms involving broader cellular rewiring in the context of established RB1 mutations rather than acute RB1 loss.

**Figure 5.**
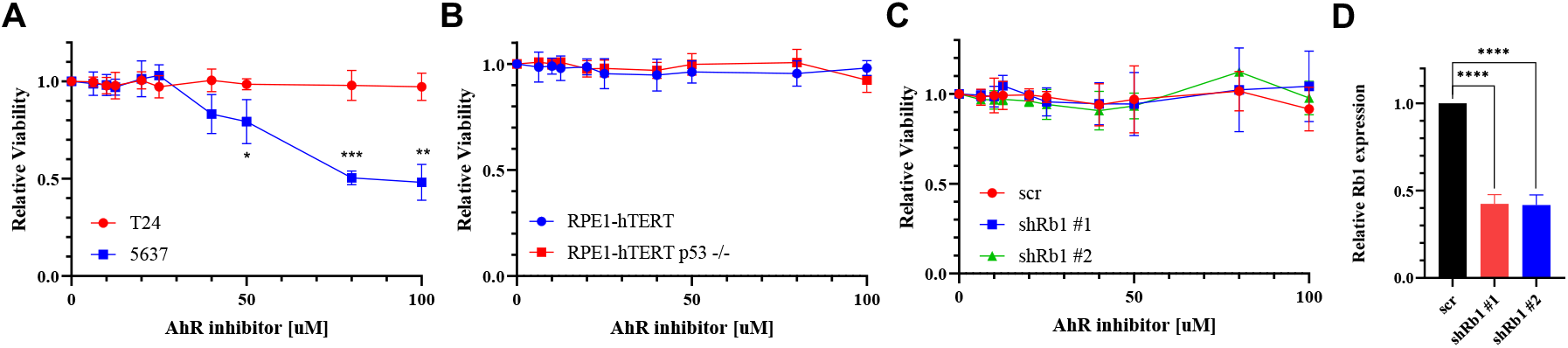
Synthetic lethality between RB1 and AhR in bladder cancer. **A**. Short-term cell viability assay in T24 and 5637 cells treated with increasing concentrations of BAY-2416964. Cell viability was measured 24 hr after drug treatment. Data are presented as mean ± s.d. (n=3). *P < 0.05;**P < 0.01;***P < 0.001 . **B**. Short-term cell viability assay in p53 WT and p53 null RPE1-hTERT cells treated with increasing concentrations of BAY-2416964. Cell viability was measured 24 hr after drug treatment. Data are presented as mean ± s.d. (n=3). **C**. Short-term cell viability assay in T24 cells expressing either scramble shRNA or shRNA targeting Rb1 and treated with increasing concentrations of BAY-2416964. Cell viability was measured 72 hr after drug treatment. Data are presented as mean ± s.d. (n=3). **D**. Quantitative RT-qPCR analysis of Rb1 knockdown in indicated T24 cell lines. ****P < 0.0001.

## Discussion

In this study, we developed SLAYER, an integrated computational framework that systematically mines multiomics data from the DepMap cancer cell line resource to nominate and prioritize synthetic lethal interactions associated with recurrent driver mutations across cancer types. Our approach combines complementary techniques examining mutational patterns alongside CRISPR gene essentiality profiles and pathway enrichment signatures. This integration enables detection of both direct genetic dependencies and indirect relationships mediated through cellular networks, providing a more comprehensive view of potential therapeutic vulnerabilities.

The systematic validation of SLAYER predictions against STRING database interactions demonstrates the biological relevance of our computational approach. The 15-fold enrichment of known protein associations among predicted synthetic lethal pairs suggests our methodology effectively captures meaningful cellular relationships rather than statistical artifacts. This enrichment is particularly noteworthy given that STRING interactions represent a conservative validation set, as many genuine synthetic lethal relationships may involve genes without direct physical interactions.

Among the high-confidence predictions, the synthetic lethal relationship between RB1 mutations and AHR inhibition in bladder cancer emerged as particularly promising. Our experimental studies confirmed increased sensitivity to AHR inhibition in RB1-mutant versus wild-type bladder cancer cells, while sparing non-cancerous cells. However, the lack of sensitization in RB1-depleted cells suggests this synthetic lethality likely arises through indirect mechanisms involving broader cellular adaptations to established RB1 loss. This observation highlights the importance of considering cellular context and adaptive responses when interpreting synthetic lethal interactions, particularly in the setting of tumor suppressor loss.

The apparent indirectness of the RB1-AHR relationship also highlights the value of our pathway-oriented analytical approach. Traditional methods focused solely on direct genetic interactions might miss such context-dependent vulnerabilities. By incorporating pathway-level information and allowing for network-mediated relationships, SLAYER can potentially identify therapeutic opportunities that emerge from complex cellular rewiring during tumor evolution. This capability is especially relevant for tumor suppressors like RB1, where acute gene loss may not fully recapitulate the cellular state that develops through prolonged adaptation to mutation.

Several limitations of our study warrant discussion. First, while our computational predictions showed strong statistical enrichment for known interactions in STRING-db, broader validation against experimental synthetic lethal datasets [**? ? ?** ] would strengthen confidence in our predictions. Second, direct comparison with other computational prediction methods would help establish SLAYER’s relative performance. These important validations were beyond the scope of the current study but represent valuable directions for future work.

Several limitations of our study warrant discussion. First, while our computational predictions showed strong statistical enrichment for known interactions, the validation experiments were restricted to a small number of cell line models. Broader experimental validation across additional cell lines and orthogonal approaches would strengthen confidence in specific predictions. Second, our current framework primarily leverages static genomic and transcriptomic profiles, potentially missing dynamic aspects of synthetic lethal relationships. Integration of temporal data or perturbation responses could provide additional insights. Third, the reliance on cancer cell lines may not fully capture the complexity of tumor biology, including interactions with the microenvironment and immune system.

Despite these limitations, SLAYER represents a valuable framework for systematic exploration of synthetic lethal interactions in cancer. The successful prediction and preliminary validation of RB1-AHR synthetic lethality demonstrates the potential for computational approaches to guide therapeutic discovery. Our comprehensive characterization of potential synthetic lethal interactions across multiple cancer types provides a rich resource for future investigation. The pathway-oriented analytical approach may be particularly valuable for understanding complex dependencies that emerge through cellular adaptation to oncogenic mutations.

Looking forward, several promising directions emerge for extending this work. Integration of additional data modalities, such as metabolomic or proteomic profiles, could provide deeper mechanistic insights. Incorporation of patient-derived models or clinical response data could help bridge the gap between cell line predictions and therapeutic applications. Finally, adaptation of our framework to analyze temporal dynamics or cellular plasticity might reveal context-dependent synthetic lethal relationships that could inform treatment strategies.

In conclusion, SLAYER provides a systematic approach for discovering potential synthetic lethal interactions with direct therapeutic relevance. While additional validation studies are warranted, our computational framework offers a valuable resource for prioritizing experimental investigation of cancer-specific vulnerabilities. The identification of AHR inhibition as a potential therapeutic strategy in RB1-mutant bladder cancer exemplifies the capacity of integrated computational analysis to reveal promising therapeutic opportunities.

## Methods

### Data Processing and Quality Control

This study utilized multi-omics data from the Cancer Dependency Map (DepMap, https://depmap.org/portal/) portal, comprising 808 cancer cell lines spanning 31 tissue types (DepMap 21Q1 release). We obtained gene expression profiles (RNA-seq), mutation annotations (DNA-seq), copy number alterations, and gene dependency scores from genome-scale CRISPR-Cas9 essentiality screens. To minimize confounding effects from genomic instability, we excluded 234 cell lines with excessive mutational burden (¿100 putative damaging mutations).

### Driver Gene-Oriented Analysis

The identification of driver genes began with characterization of non-silent mutations (missense, nonsense, frameshift, or splice site) across the cell line panel. We defined recurrent mutations as those occurring at ≥ 5% frequency either across all cell lines (pan-cancer analysis) or within specific cancer types (tissue-specific analysis). Each mutation’s functional impact was assessed using established computational tools including SIFT and PolyPhen, with additional annotation from cancer gene databases such as COSMIC and OncoKB.

For each identified driver gene, we stratified cell lines into mutant and wild-type groups, requiring a minimum of 3 cell lines per group for statistical validity. We then calculated the delta dependency (DD) score between groups as:

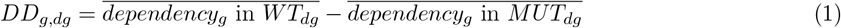

where *g* represents the gene being tested and *dg* represents the driver gene. Statistical significance was assessed using two-sample t-tests with Benjamini-Hochberg correction for multiple testing. Synthetic lethal interactions were identified using thresholds of adjusted p-value ¡ 0.05 and DD ¿ 0.2.

### Pathway Enrichment-Oriented Analysis

Our pathway-level analysis framework incorporated 189 canonical oncogenic pathway signatures from the MSigDB C6 collection, focusing on experimentally validated gene sets reflecting oncogenic states. We quantified pathway activity using single-sample Gene Set Enrichment Analysis (ssGSEA) and normalized enrichment scores across cell lines. These scores were then correlated with gene dependency profiles to construct a weighted bipartite graph connecting genes and pathways.

Edge weights in the network were determined by correlation coefficients, with edges filtered using an absolute correlation threshold of 0.5. The network visualization employed a sigmoid-based transparency function:

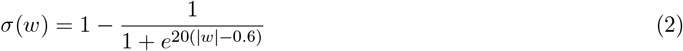

To identify potential pathway-mediated relationships, we implemented a Breadth-First Search algorithm to discover connecting paths between driver gene mutations, gene dependencies, and intermediate pathway nodes. Path scoring incorporated both edge weights and directionality of correlations.

### Candidate SLI Prioritization

The prioritization of synthetic lethal interactions followed a systematic framework integrating multiple evidence sources. Statistical significance requirements included adjusted p-value ¡ 0.05 and DD ¿ 0.2 across both analytical approaches. Clinical relevance was assessed through TCGA mutation frequency data from cBioPortal, requiring ¿15% frequency in the relevant cancer type. Therapeutic potential was evaluated using the DepMap PRISM repurposing database to identify druggable target genes.

We validated predicted interactions through comparison with the STRING database and calculated enrichment versus random gene pairs. Statistical analyses were performed using R (version 4.1.0), with multiple testing correction applied both within each cancer type and across the pan-cancer analysis. All code and detailed documentation are available at https://github.com/zivc888/SLICE.

### Experimental validation of the SLI between RB1 and AhR in bladder cancer

#### Cell lines and cell culture

T24 and 5637 cell lines were grown in RPMI-1640 medium (Gibco) supplemented with 10% heat-inactivated FBS, 2 mM L-glutamine and 100 mg/ml penicillin/streptomycin. RPE1-hTERT wild-type and p53 -/- cell lines were grown in Dulbecco’s modified Eagle’s medium (Gibco) supplemented with 10% heat-inactivated FBS, 2 mM L-glutamine and 100 mg/ml penicillin/streptomycin.

#### Lentiviral transduction

Lentiviral particles were generated by co-transfecting HEK293T cells plated in 10cm plates with 1.64pmol target vector, together with 1.3pmol psPAX2 (Addgene #12260) and 0.72pmol pMD2.G (Addgene #12259) using 3:1 *µ*g PEI to *µ*g DNA ratio. Media containing the viral particles were collected 48 hr post-transfection and filtered with 0.45 *µ*m filters. The indicated cell lines were transduced with the lentiviral particles in the presence of 10*µ*g/ml polybrene (Sigma-Aldrich H9268). At 48 hr post-infection, transduced cells were selected using the appropriate antibiotics.

#### Short hairpin RNA (shRNA) knockdown

shRNA oligonucleotides directed against RB1 were annealed and cloned into pLKO-puro lentiviral vector (Addgene #10878) digested with EcoRI and AgeI. Lentiviral particles were generated as described above and used to transduce T24 cells, followed by selection with 1*µ*g/ml puromycin (Invivogen #ant-pr) for 3 days. Cells were maintained in complete medium in the presence of 0.6*µ*g/ml puromycin. Gene expression knockdown was validated by RT-qPCR. The shRNA and qPCR primer sequences used in this study are included in Supplementary Table 2.

#### RNA isolation and qPCR

Total RNA was isolated from cells using the TRIzol reagent according to the manufacturer’s instructions (Sigma). 1*µ*g RNA was used for cDNA synthesis using the qScript cDNA Synthesis Kit (Quanta) with random primers. Real-time qPCR for measuring mRNA levels was performed using Step-One-Plus real-time PCR System (Applied Biosystems) and the Fast SYBR Green Master mix (Applied Biosystems) with three technical repeats for each PCR. Data analysis and quantification were performed using StepOne software V2.2 supplied by Applied Biosystems. Primer sequences used for qPCR are included in Supplementary Table 2.

#### AhR inhibitor

BAY-2416964 was purchased from Selleckchem.

#### Short-term growth delay assay

For determining drug sensitivity, cells were seeded in 96-well plates in duplicates at a density of 3,000-7,500 cells per well. 24 hr post-seeding, BAY-2416964 was added at the indicated concentrations. Cell viability was measured at the indicated timepoints using the CellTiter 96® AQueous One Solution Cell Proliferation Assay (Promega) following the manufacturer’s protocol, and absorbance was measured using Epoch Microplate Spectrophotometer (BioTek). Cell viability was normalized to the viability of untreated cells.

## Declarations

## Data and code availability

All data used in this study is publicly available at the DepMap portal. The code developed in this study is available at https://github.com/zivc888/SLICE.

## Authors Contribution

The authors confirm their contribution to the paper as follows: study conception and design: DA; algorithm development: ZC; analysis and interpretation of results: ZC & DA; experimental validation: CYBU, ASB, EAZ & NA; draft manuscript preparation: ZC, EP & DA. All authors reviewed the results and approved the final version of the manuscript.

## Acknowledgments

We thank Hinanit Kolati for generously sharing cell lines. DA is supported by the Azrieli Faculty Fellowship. ASB is supported by Joan and Irwin Jacobs fellowship. NA is supported by the Neubauer Family foundation. We thank the Aran Lab members for helpful comments.

