## Supplementary Figures for "SLAYER: A Computational Framework for Identifying Synthetic Lethal Interactions through Integrated Analysis of Cancer Dependencies"

### Supplementary Figure 1

Here we present heatmaps depicting the top synthetic lethal interaction (SLI) pairs in each cancer type (similar to Figure 4A), with redder colors indicating higher delta dependency scores.

#### Bladder cancer

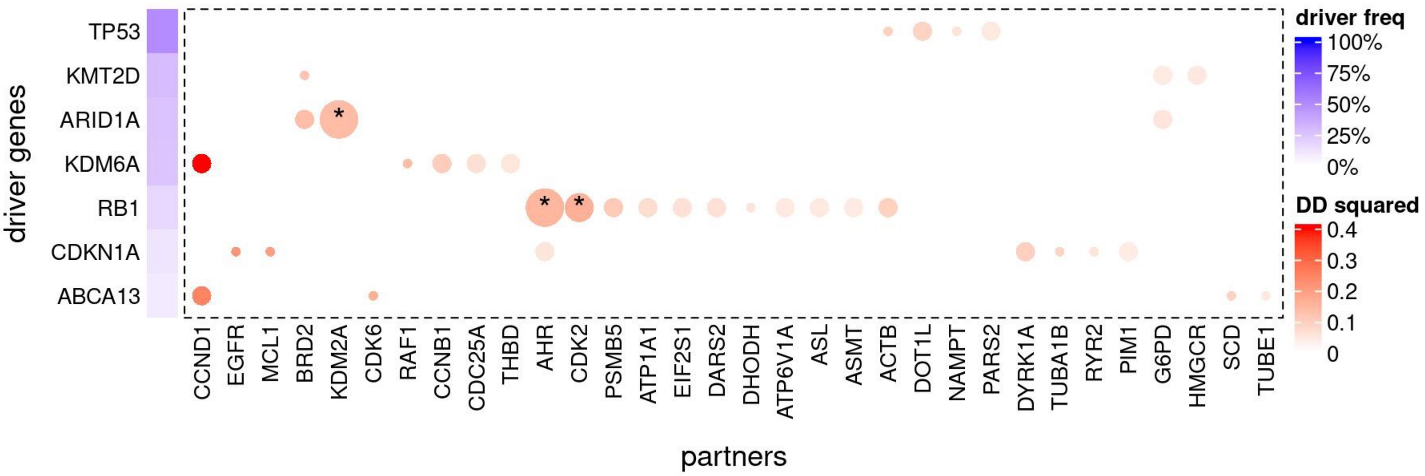


#### Brain cancer

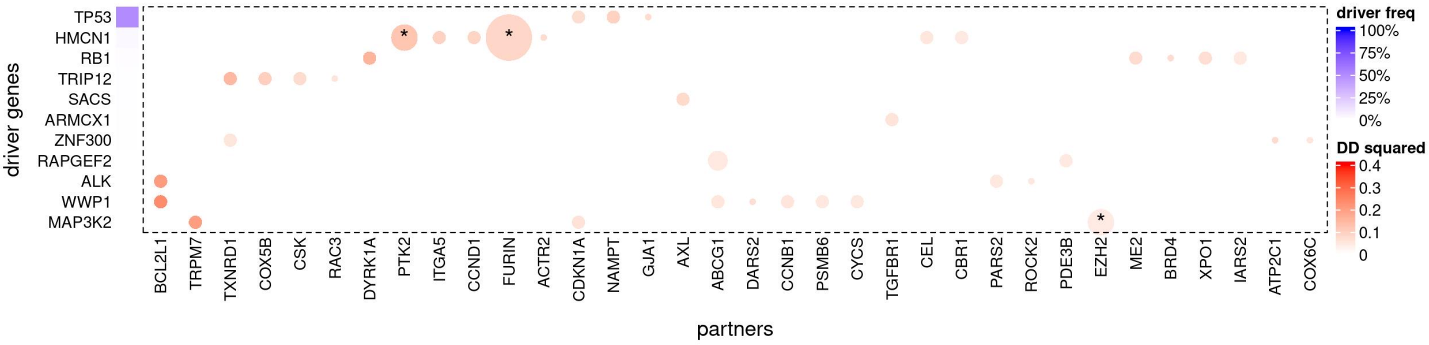


#### Breast cancer

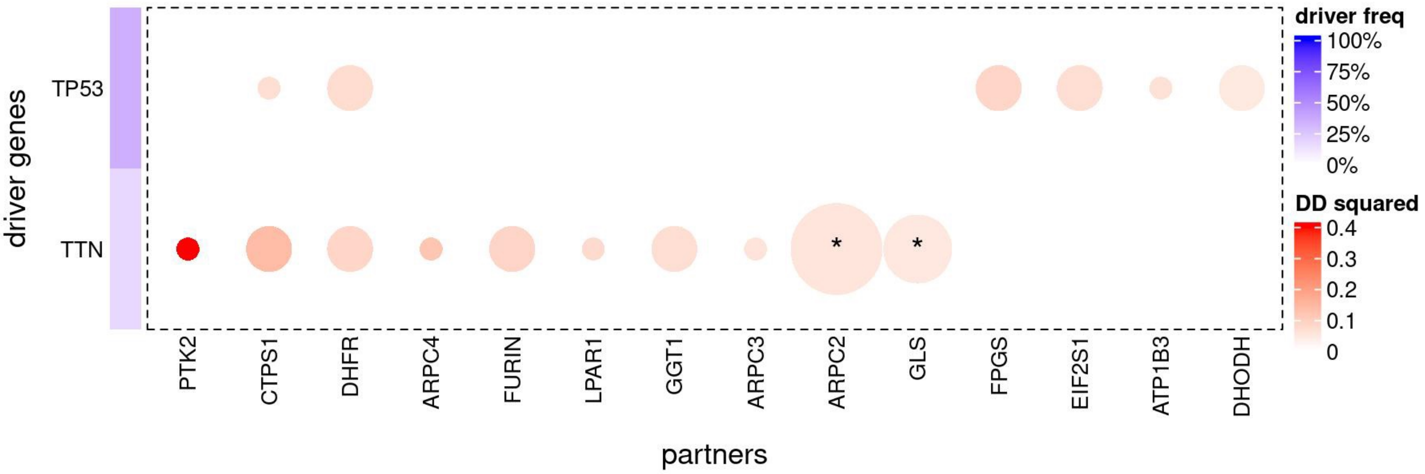

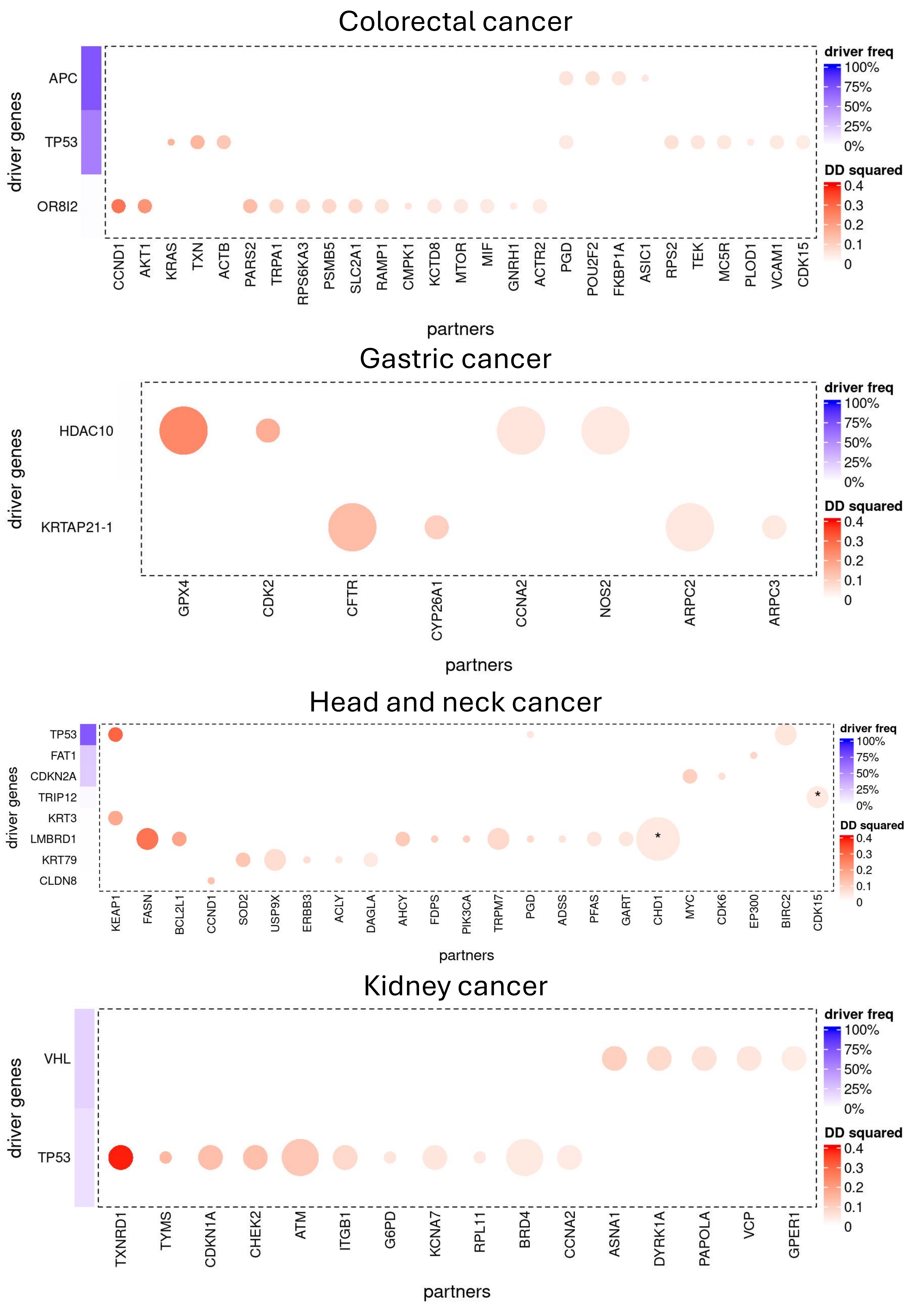
Colorectal cancer
driver genes
APC
TP53
OR8I2
CCND1
AKT1
KRAS
TXN
ACTB
PARS2
TRPA1
RPS6KA3
PSMB5
SLC2A1
RAMP1
CMPK1
KCTD8
MTOR
MIF
GNRH1
ACTR2
PGD
POU2F2
FKBP1A
ASIC1
RPS2
TEK
MC5R
PLOD1
VCAM1
CDK15
partners
driver freq
100%
75%
50%
25%
0%
DD squared
0.4
0.3
0.2
0.1
0
Gastric cancer
driver genes
HDAC10
KRTAP21-1
GPX4
CDK2
CFTR
CYP26A1
CCNA2
NOS2
ARPC2
ARPC3
partners
driver freq
100%
75%
50%
25%
0%
DD squared
0.4
0.3
0.2
0.1
0
Head and neck cancer
driver genes
TP53
FAT1
CDKN2A
TRIP12
KRT3
LMBRD1
KRT79
CLDN8
KEAP1
FASN
BCL2L1
CCND1
SOD2
USP9X
ERBB3
ACLY
DAGLA
AHCY
FDPS
PIK3CA
TRPM7
PGD
ADSS
PFAS
GART
CHD1
MYC
CDK6
EP300
BIRC2
CDK15
*
*
partners
driver freq
100%
75%
50%
25%
0%
DD squared
0.4
0.3
0.2
0.1
0
Kidney cancer
driver genes
VHL
TP53
TXNRD1
TYMS
CDKN1A
CHEK2
ATM
ITGB1
G6PD
KCNA7
RPL11
BRD4
CCNA2
ASNA1
DYRK1A
PAPOLA
VCP
GPER1
partners
driver freq
100%
75%
50%
25%
0%
DD squared
0.4
0.3
0.2
0.1
0

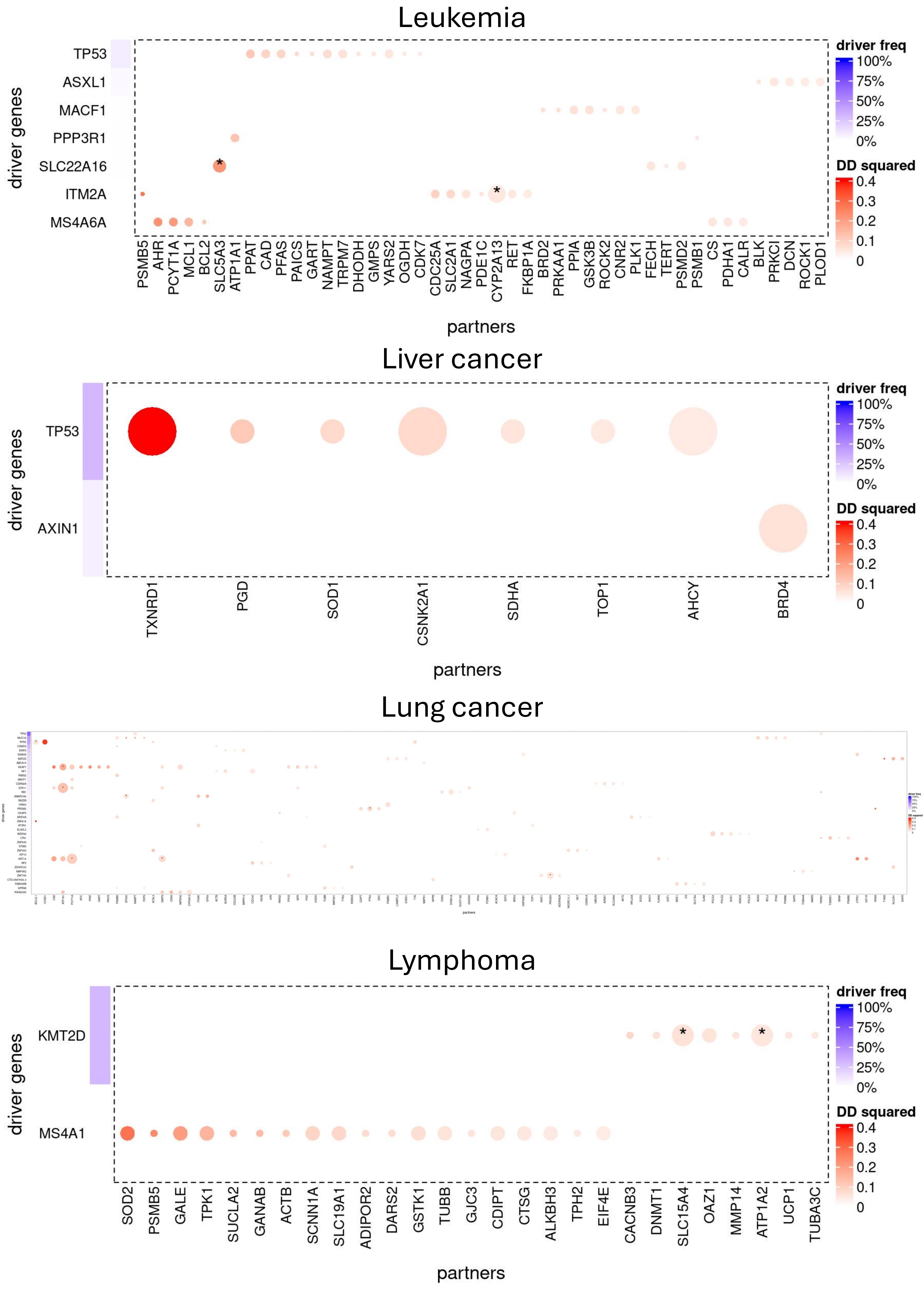
Leukemia
driver genes
TP53
ASXL1
MACF1
PPP3R1
SLC22A16
ITM2A
MS4A6A
PSMB5 AHR PCYT1A MCL1 BCL2 SLC5A3 ATP1A1 PPAT CAD PFAS PAICS GART NAMPT TRPM7 DHODH GMPS YARS2 OGDH CDK7 CDC25A SLC2A1 NAGPA PDE1C CYP2A13 RET FKBP1A BRD2 PRKAA1 PPIA GSK3B ROCK2 CNR2 PLK1 FECH TERT PSMD2 PSMB1 CS PDHA1 CALR BLK PRKCI DCN ROCK1 PLOD1
partners
driver freq
100%
75%
50%
25%
0%
DD squared
0.4
0.3
0.2
0.1
0
Liver cancer
driver genes
TP53
AXIN1
TXNRD1 PGD SOD1 CSNK2A1 SDHA TOP1 AHCY BRD4
partners
driver freq
100%
75%
50%
25%
0%
DD squared
0.4
0.3
0.2
0.1
0
Lung cancer
Lymphoma
driver genes
KMT2D
MS4A1
SOD2 PSMB5 GALE TPK1 SUCLA2 GANAB ACTB SCNN1A SLC19A1 ADIPOR2 DARS2 GSTK1 TUBB GJC3 CDIPT CTSG ALKBH3 TPH2 EIF4E CACNB3 DNMT1 SLC15A4 OAZ1 MMP14 ATP1A2 UCP1 TUBA3C
partners
driver freq
100%
75%
50%
25%
0%
DD squared
0.4
0.3
0.2
0.1
0

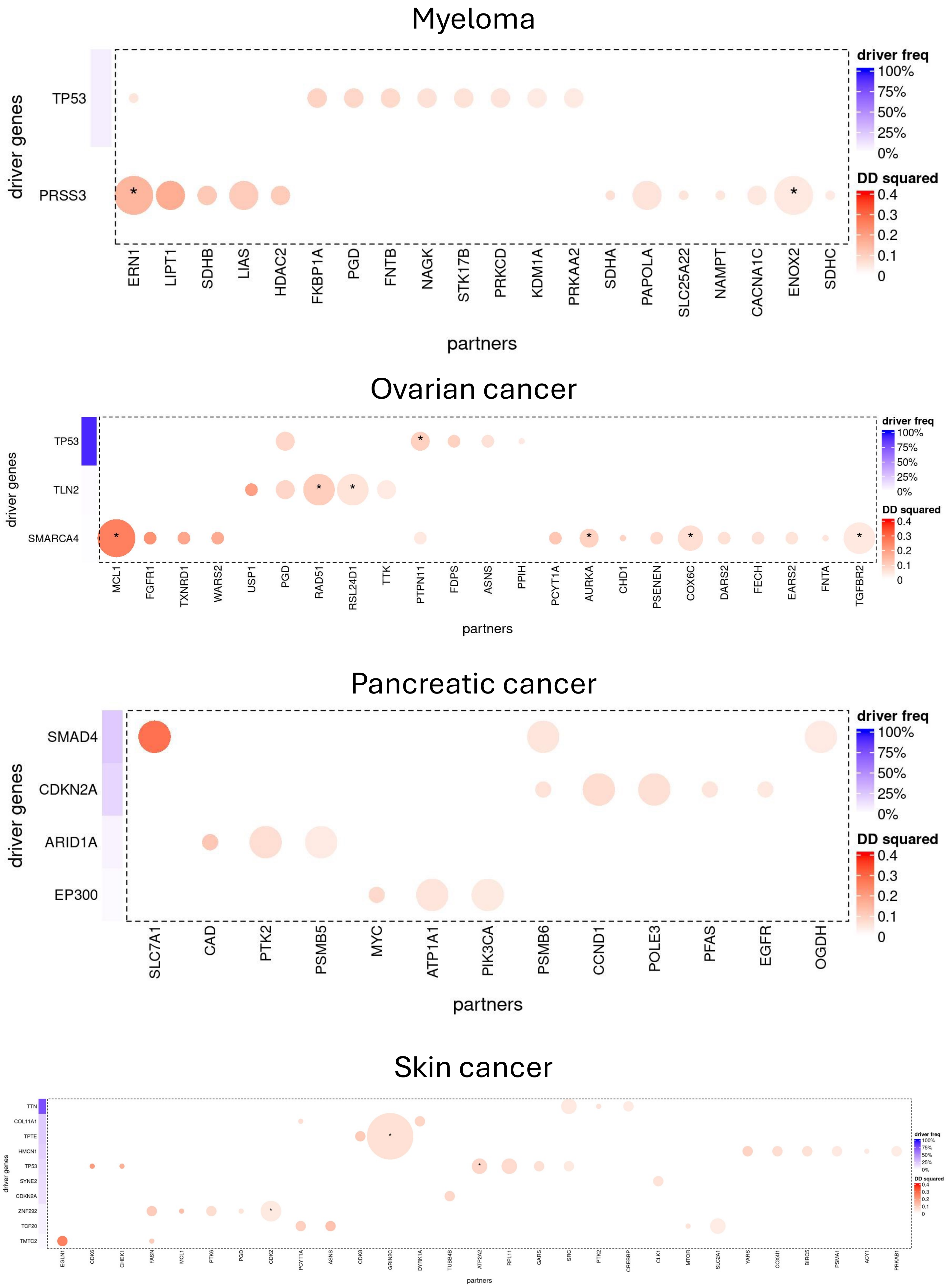

Myeloma
driver genes
TP53
PRSS3
partners
ERN1
LIPT1
SDHB
LIAS
HDAC2
FKBP1A
PGD
FNTB
NAGK
STK17B
PRKCD
KDM1A
PRKAA2
SDHA
PAPOLA
SLC25A22
NAMPT
CACNA1C
ENOX2
SDHC
driver freq
100%
75%
50%
25%
0%
DD squared
0.4
0.3
0.2
0.1
0
Ovarian cancer
driver genes
TP53
TLN2
SMARCA4
partners
MCL1
FGFR1
TXNRD1
WARS2
USP1
PGD
RAD51
RSL24D1
TTK
PTPN11
FDPS
ASNS
PPIH
PCYT1A
AURKA
CHD1
PSENEN
COX6C
DARS2
FECH
EARS2
FNTA
TGFBR2
driver freq
100%
75%
50%
25%
0%
DD squared
0.4
0.3
0.2
0.1
0
Pancreatic cancer
driver genes
SMAD4
CDKN2A
ARID1A
EP300
partners
SLC7A1
CAD
PTK2
PSMB5
MYC
ATP1A1
PIK3CA
PSMB6
CCND1
POLE3
PFAS
EGFR
OGDH
driver freq
100%
75%
50%
25%
0%
DD squared
0.4
0.3
0.2
0.1
0
Skin cancer
driver genes
TTN
COL11A1
TPTE
HMCN1
TP53
SYNE2
CDKN2A
ZNF292
TCF20
TMTC2
partners
EGLN1
CDK6
CHEK1
FASN
MCL1
PTK6
PGD
CDK2
PCYT1A
ASNS
CDK8
GRIN2C
DYRK1A
TUBB4B
ATP2A2
RPL11
GARS
SRC
PTK2
CREBBP
CLK1
MTOR
SLC2A1
YARS
COX4I1
BIRC5
PSMA1
ACY1
PRKAB1
driver freq
100%
75%
50%
25%
0%
DD squared
0.4
0.3
0.2
0.1
0

### Supplementary Figure 2

Here we present plots for some of the top potential synthetic lethal interaction (SLI) pairs (similar to Figure 4B-D). A) Volcano plot showing all potential synthetic lethal interactions in different cancer types for different mutations. B) Boxplots illustrating the separation in gene dependency scores for the top pair for the corresponding mutation. C) Dependency networks for the corresponding paired gene showing the association with the mutation pathway.

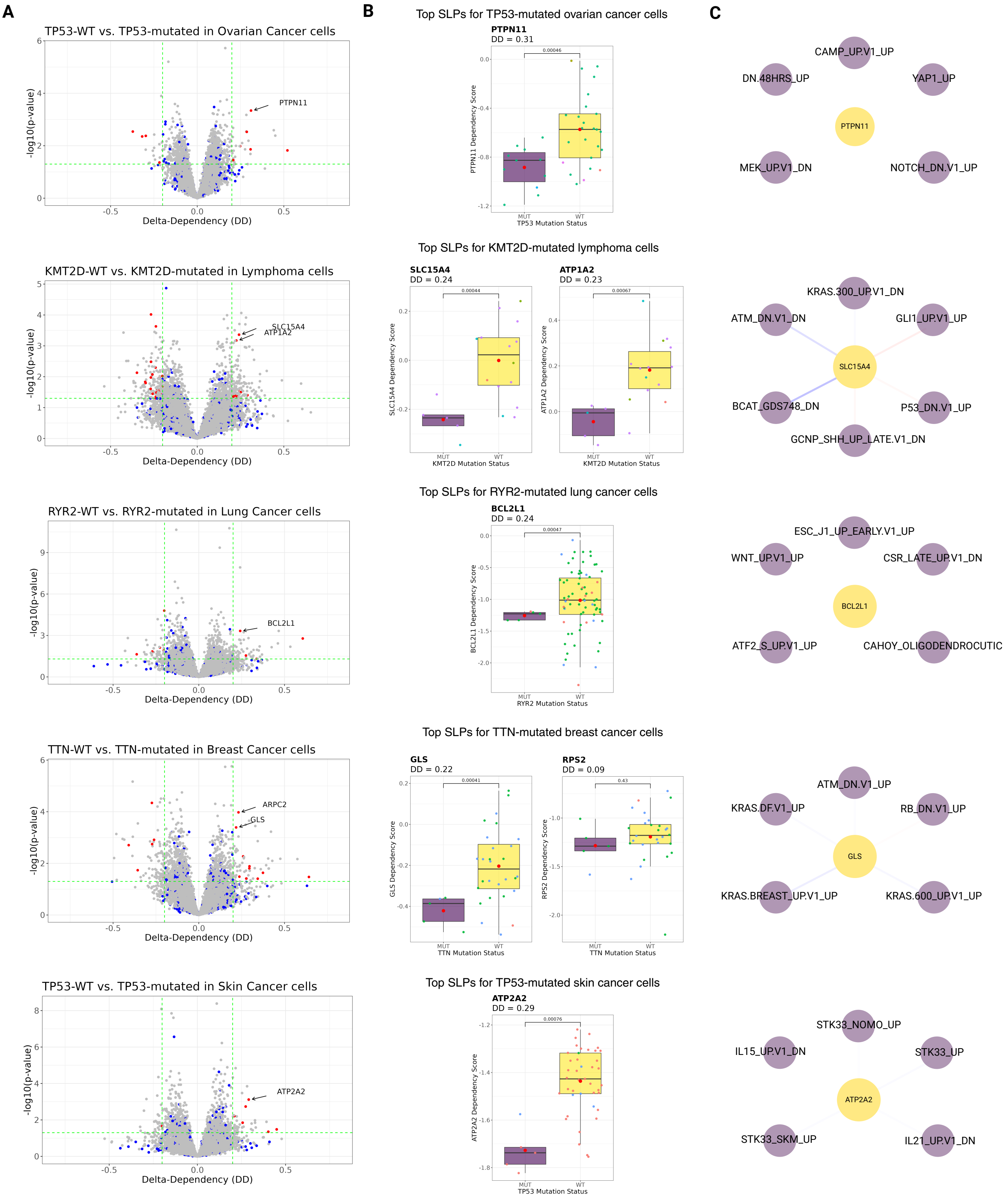
